## Supplemental Figure 1-1 for "GAF is essential for zygotic genome activation and chromatin accessibility in the early *Drosophila* embryo"

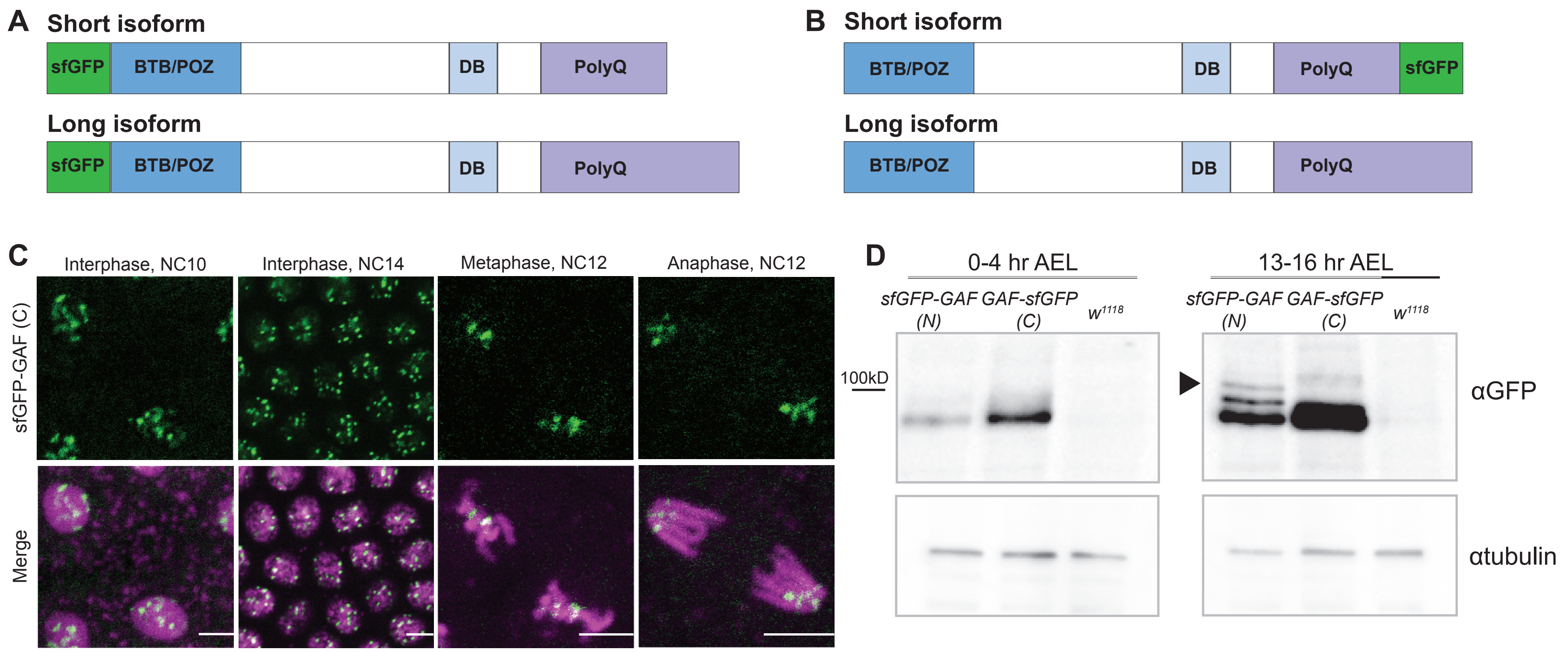


**Figure 1- figure supplement 1: N- and C-terminal GFP-tags label distinct GAF isoforms** A. Cartoon representation of the N-terminal sfGFP tag on both GAF protein isoforms B. Cartoon representation of C-terminal sfGFP tag on the short GAF protein isoform. C. Confocal images of living *His2Av-RFP; GAF-sfGFP(C)* embryos in interphase during NC10 and NC14 and mitosis during NC12. His2Av-RFP is shown in magenta. sfGFP-GAF (C) is shown in green. Scale bars, 5 μm. D. Immunoblot on embryo extract from *sfGFP-GAF(N)* homozygous, *GAF-sfGFP(C)* homozygous, and $w^{1118}$ lines 0-4 hour after egg laying (AEL) and 13-16 hours AEL with an anti-GFP antibody. αtubulin is shown as a loading control. Arrowhead indicates the long GAF isoform.
