## Supplemental Figure 1-2 for "GAF is essential for zygotic genome activation and chromatin accessibility in the early *Drosophila* embryo"

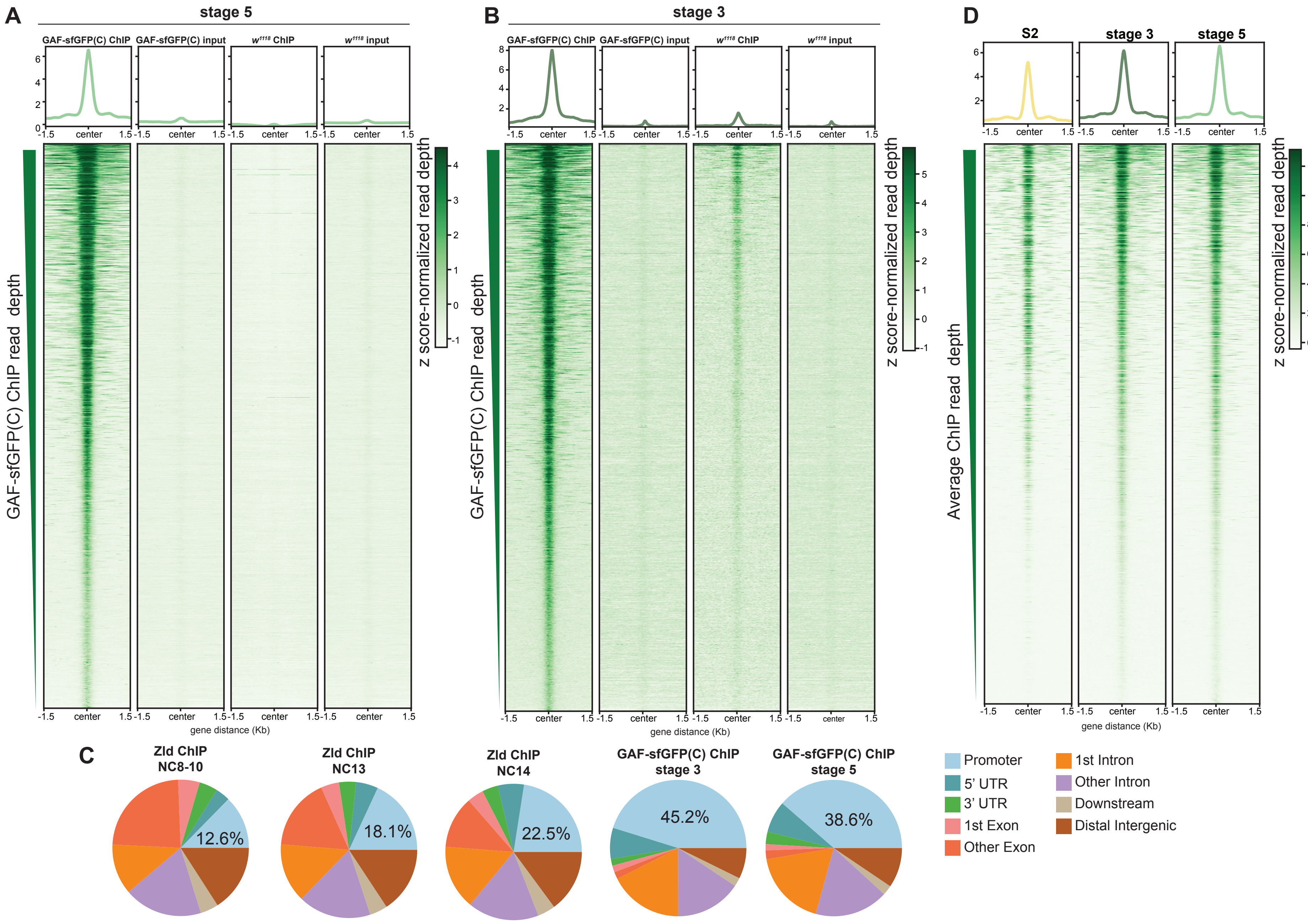


**Figure 1 - figure supplement 2: GAF binds thousands of regions in the embryo and S2 cells.** A. Heatmap of stage 5 GAF-sfGFP(C) ChIP, GAF-sfGFP(C) input, *w1118* ChIP, and *w1118* input peaks (n = 4175). Immunoprecipitation was performed with an anti-GFP antibody. B. Heatmap of stage 3 GAF-sfGFP(C) ChIP, GAF-sfGFP(C) input, *w1118* ChIP, and *w1118* input peaks (n = 3391). Immunoprecipitation was performed with an anti-GFP antibody. C. Genomic distribution of Zld binding at NC8-10, NC13, and NC14 (Harrison et al. 2011) and GAF binding at stage 3 and stage 5. D. Heatmaps of GAF binding in S2 cells (Fuda et al. 2015) and GAF-sfGFP(C) binding in stage 3 and stage 5 embryos for all 4175 regions with a stage 5 ChIP peak.
