## Supplemental Figure 1-3 for "GAF is essential for zygotic genome activation and chromatin accessibility in the early *Drosophila* embryo"

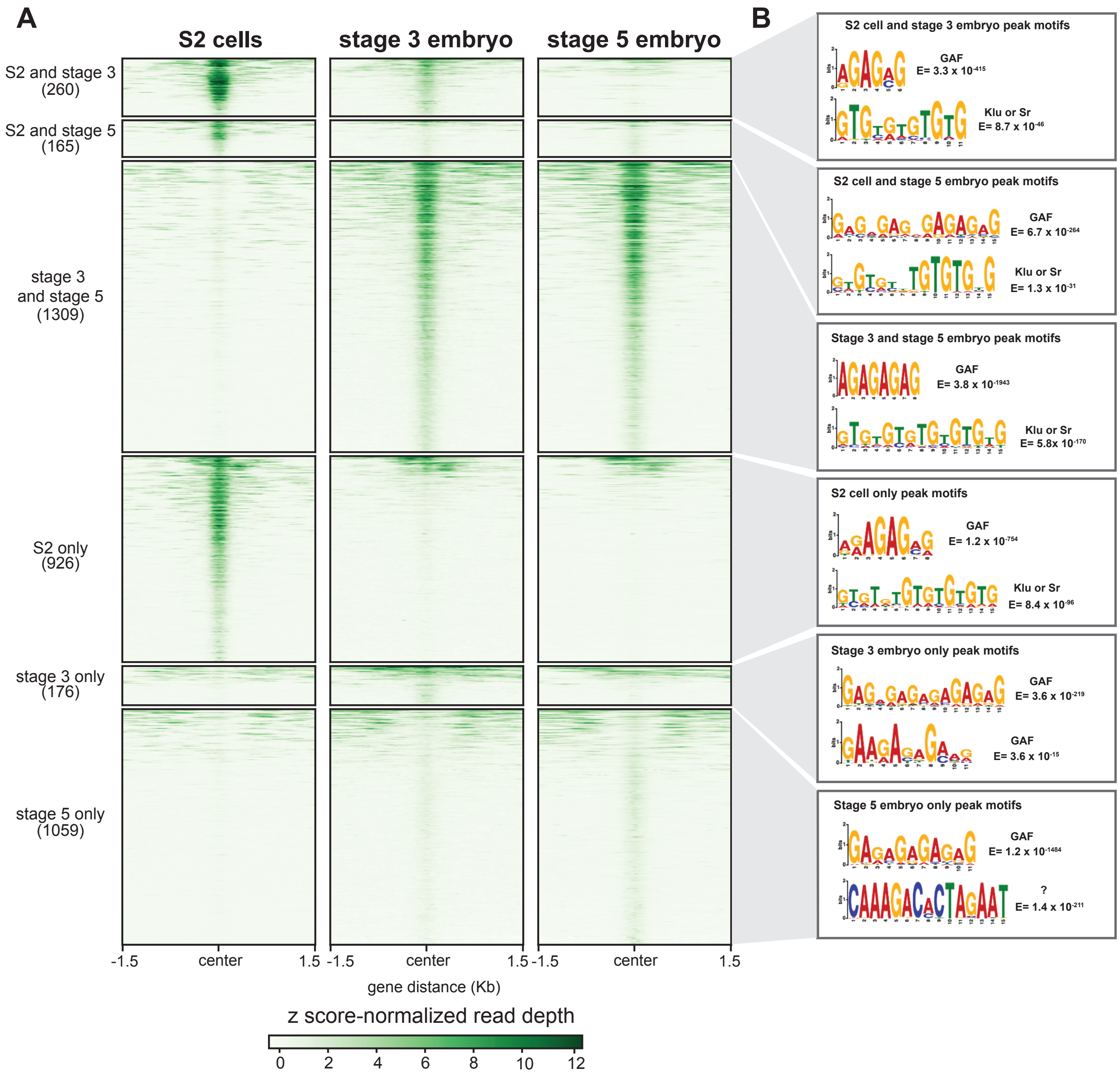


**Figure 1 - figure supplement 3- GAF has tissue-specific binding.** A. Heatmap of GAF ChIP-seq peaks in S2 cells (Fuda et al. 2015) and GAF-sfGFP(C) ChIP-seq peaks from stage 3 and stage 5 embryos (this work), excluding peaks that are shared in all three datasets. The heatmap is divided into subclasses as labelled. B. The top two motifs enriched in each subclass of peaks as identified by MEME suite. Note the top motif in each subclass is a GA-rich motif known to be bound by GAF.
