## Supplemental Figure 2-1 for "GAF is essential for zygotic genome activation and chromatin accessibility in the early *Drosophila* embryo"

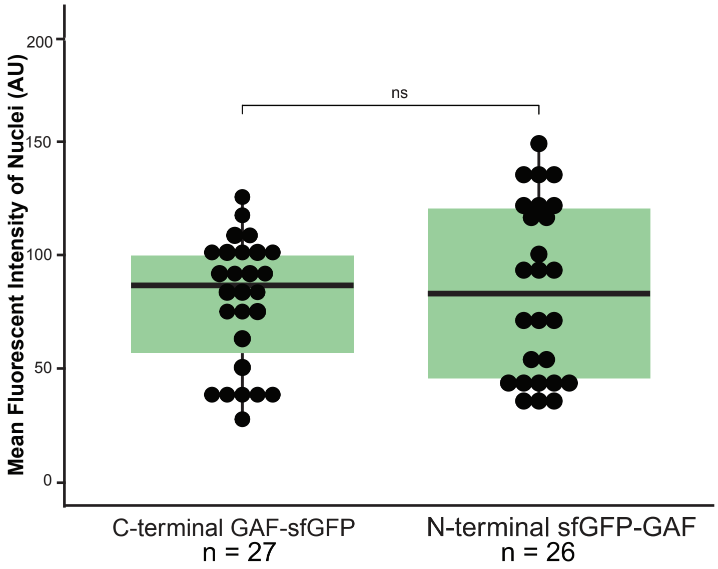


**Figure 2 - figure supplement 1: sfGFP-GAF(N) and GAF-sfGFP(C) are expressed at similar levels in the early embryo.** Boxplots showing the mean fluorescent intensity (arbitrary units, AU) of 10 nuclei in *GAF-sfGFP (C)* homozygous embryos (n=27) and *sfGFP-GAF (N)* homozygous embryos (n=26). The box indicates the lower (25%) and upper (75%) quantiles, and the solid line indicates the median. p value = 0.5192 as determined by Wilcoxon rank sum test.
