## Supplemental Figure 3-1 for "GAF is essential for zygotic genome activation and chromatin accessibility in the early *Drosophila* embryo"

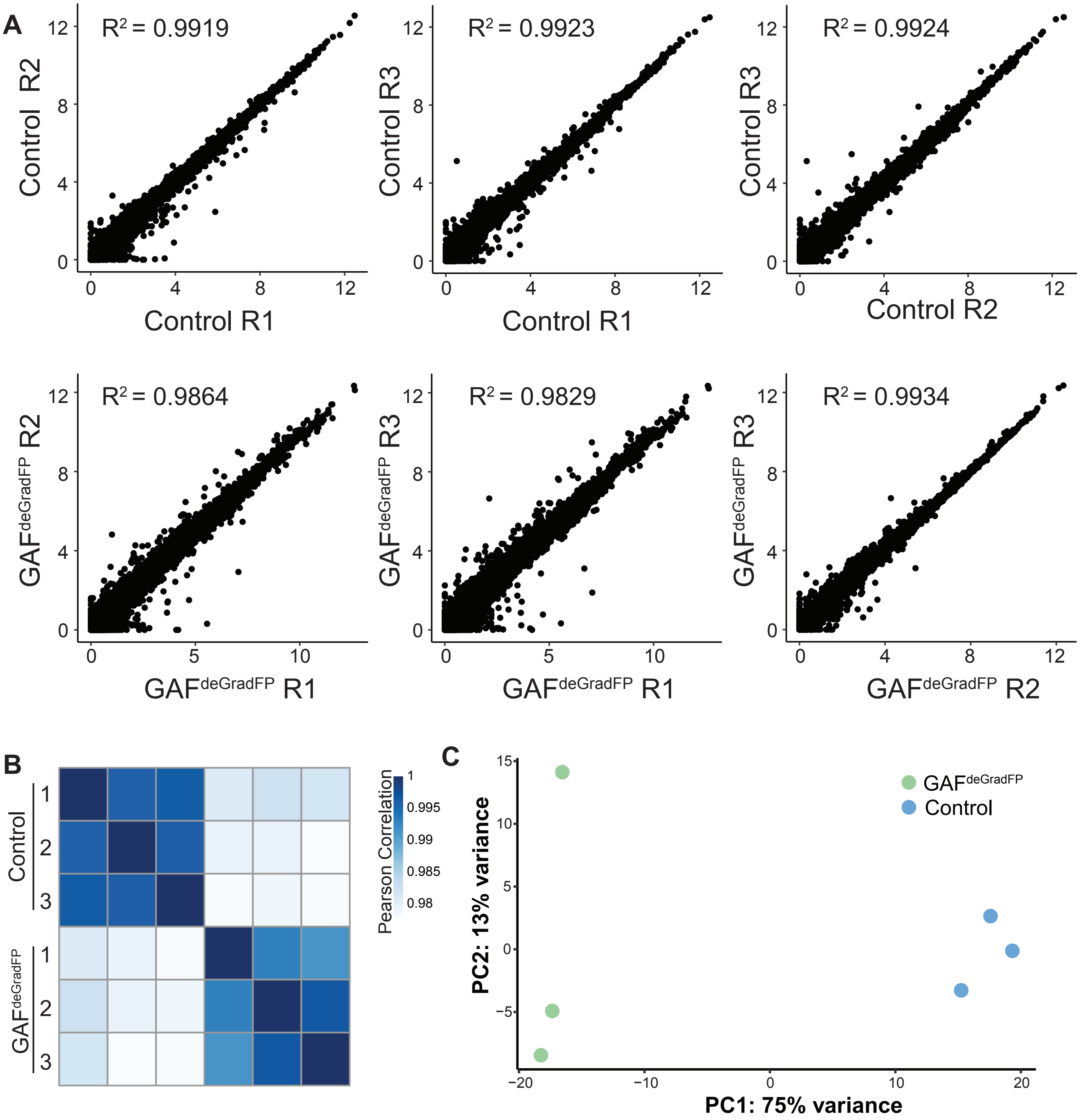


**Figure 3 - figure supplement 1: RNA-seq replicates are reproducible.** A. Pairwise scatterplots showing correlation between the three RNA-seq replicates for GAF$^{deGradFP}$ and *sfGFP-GAF(N)* homozgyous controls. Pearson correlation coefficient shown. B. Heatmap of Pearson correlation coefficient between RNA-seq replicates for GAF$^{deGradFP}$ and *sfGFP-GAF(N)* homozgyous controls. C. PCA plot of RNA-seq replicates from GAF$^{deGradFP}$ and *sfGFP-GAF(N)* homozgyous controls
