## Supplemental Figure 3-2 for "GAF is essential for zygotic genome activation and chromatin accessibility in the early *Drosophila* embryo"

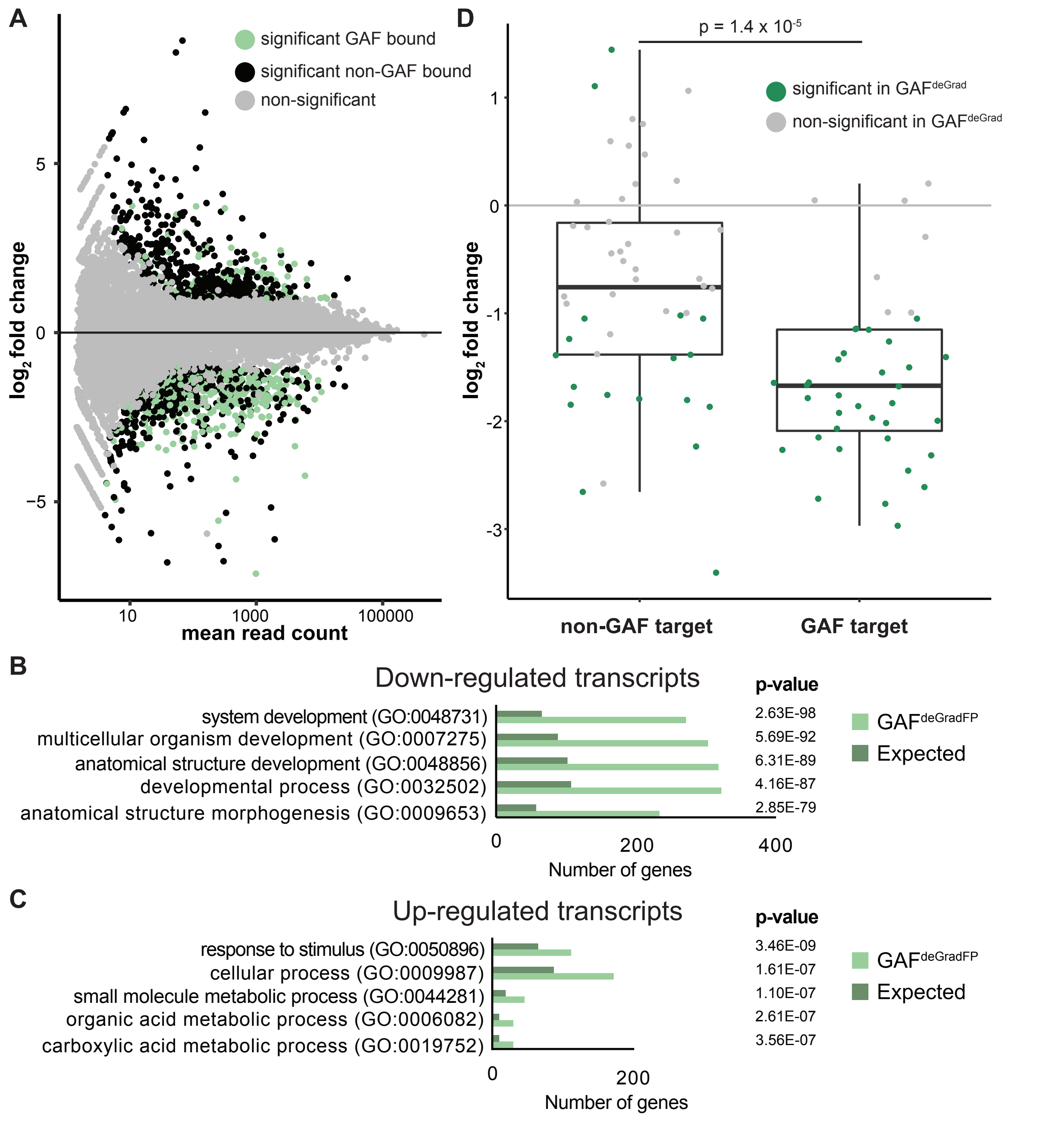


**Figure 3 - figure supplement 2: RNA-seq identifies genes mis-regulated in GAF$^{deGradFP}$ embryos.** A. MA plot of transcripts mis-expressed in GAF$^{deGradFP}$ embryos as compared to *sfGPF-GAF(N)* homozygous controls. Stage 5 GAF-sfGFP(C) ChIP-seq was used to identify GAF-bound target genes. B. Gene Ontology (GO) term analysis was performed on transcripts down-regulated in GAF$^{deGradFP}$ embryos compared to controls. C. Gene Ontology (GO) term analysis was performed on transcripts up-regulated in GAF$^{deGradFP}$ embryos compared to controls. D. $\log_2$ fold change of genes expressed at NC14 (Li et al. 2014, “later” classes) in GAF$^{deGrad}$ embryos compared to *sfGFP-GAF (N)* homozygous controls . Genes are divided by those that have a proximal GAF-binding site as identified by ChIP-seq (GAF target) and those that do not (non-GAF target). Color indicates those that are significantly changed in the GAF$^{deGrad}$ embryos as compared to *sfGFP-GAF(N)* controls. GAF-target genes are significantly more down-regulated than non-GAF target genes ($p = 1.4 \times 10^{-5}$, Wilcoxon rank sum test).
