## Supplemental Figure 3-3 for "GAF is essential for zygotic genome activation and chromatin accessibility in the early *Drosophila* embryo"

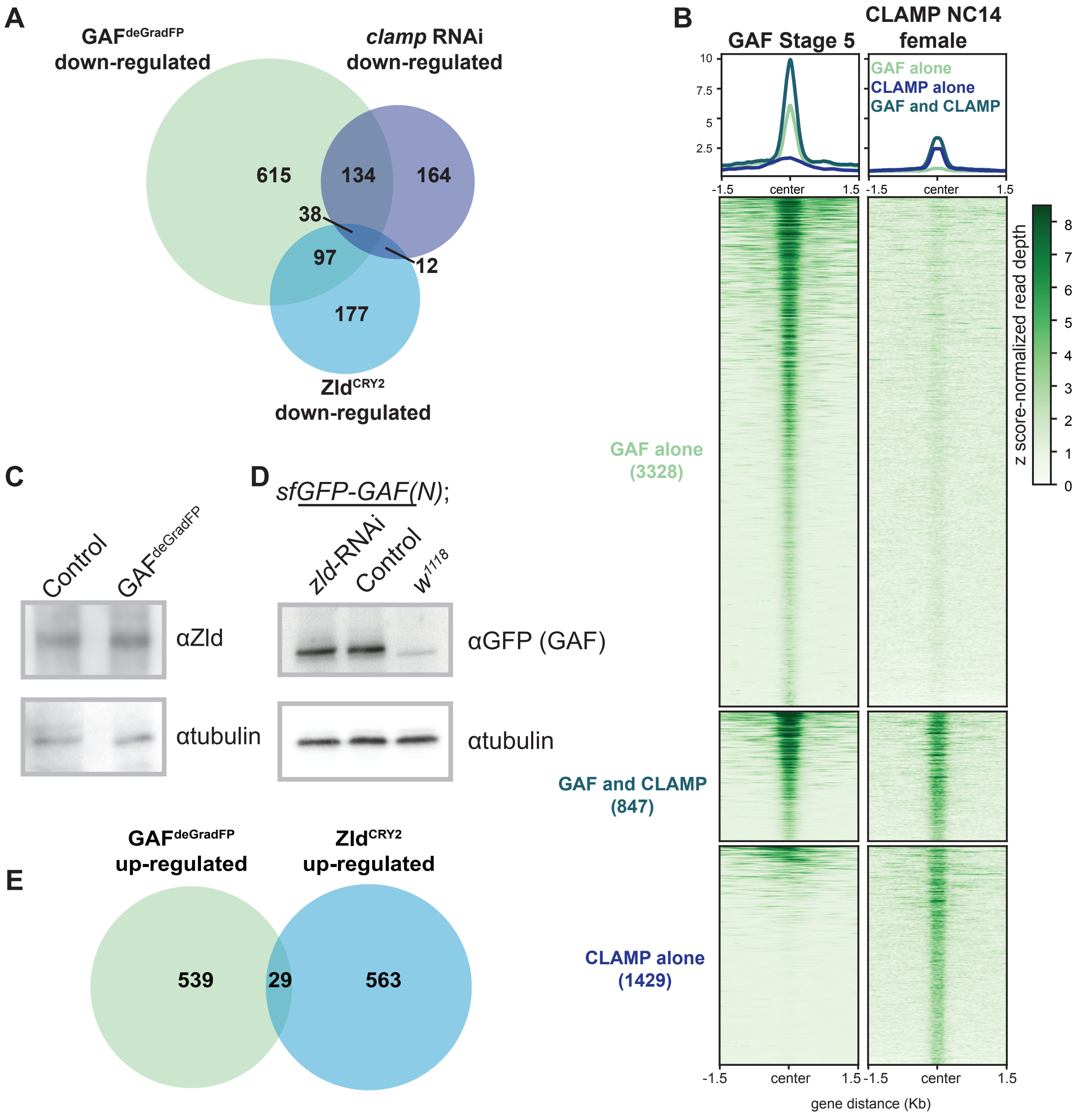


**Figure 3 - figure supplement 3: GAF regulates genes distinct from Zld and CLAMP.** A. Overlap of transcripts down-regulated in GAF$^{deGradFP}$ embryos, down-regulated in Zld$^{CRY2}$ embryos at NC14 (McDaniel et al. 2019), and down-regulated in *clamp*-RNAi 2-4 hour embryos (Rieder et al. 2017). B. Heatmap of GAF-sfGFP(C) ChIP peaks at stage 5 and CLAMP ChIP peaks in female embryos at NC14. CLAMP ChIP-seq from Reider et al. 2019. C. Immunoblot of extract from 2-2.5 hr AEL GAF$^{deGradFP}$ and *sfGFP-GAF(N)* homozygous control embryos using anti-Zld antibody. Tubulin is shown as a loading control. D. Immunoblot of extract from 2-2.5 hr AEL *sfGFP-GAF(N)* embryos in which *zld* was depleted (*zld*-RNAi) and in which *zld* levels were unperturbed (Control) probed with anti-GFP antibody. *w$^{1118}$* embryos are included as controls for anti-GFP immunoreactivity. Tubulin is shown as a loading control. E. Overlap of transcripts up-regulated in GAF$^{deGradFP}$ embryos and Zld$^{CRY2}$ embryos (McDaniel et al. 2019) ($p = 0.64$, $\log_2$(odds ratio) = -0.16, two-tailed Fisher's exact test).
