## Supplemental Figure 4-1 for "GAF is essential for zygotic genome activation and chromatin accessibility in the early *Drosophila* embryo"

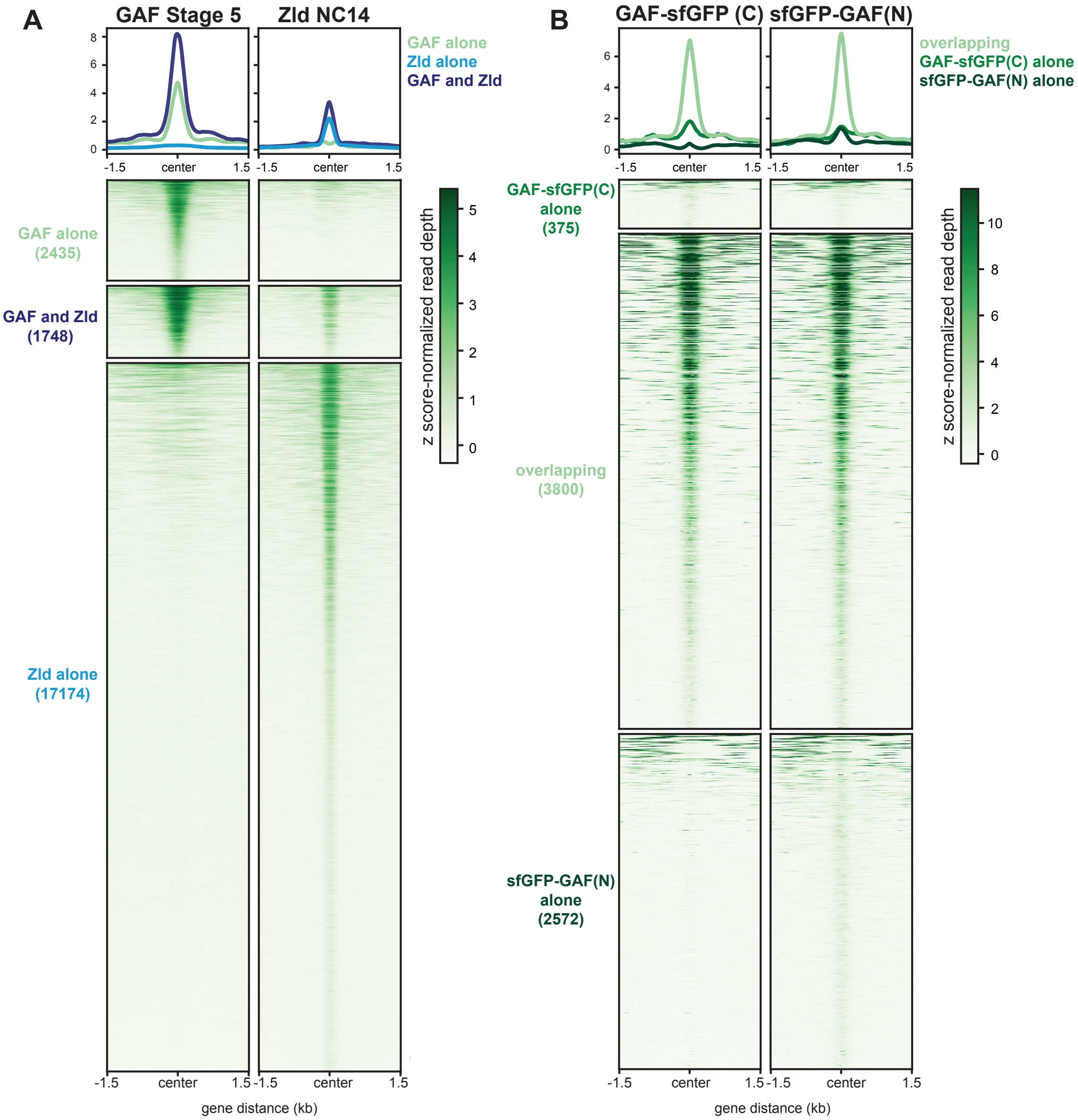


**Figure 4 - figure supplement 1: GAF and Zld bind to shared and unique regions of the genome.** A. Heatmaps of GAF-sfGFP(C) ChIP peaks at stage 5 and Zld ChIP peaks at NC14. Zld ChIP-seq from Harrison et al. 2011. B. Heatmap of anti-GFP ChIP-seq peaks from *GAF-sfGFP(C)* homozygous stage 5 embryos and *sfGFP-GAF(N)*/+ heterozygous 2-2.5 hr AEL embryos. ChIP peaks show reproducible identification of the highest peaks.
