## Supplemental Figure 4-2 for "GAF is essential for zygotic genome activation and chromatin accessibility in the early *Drosophila* embryo"

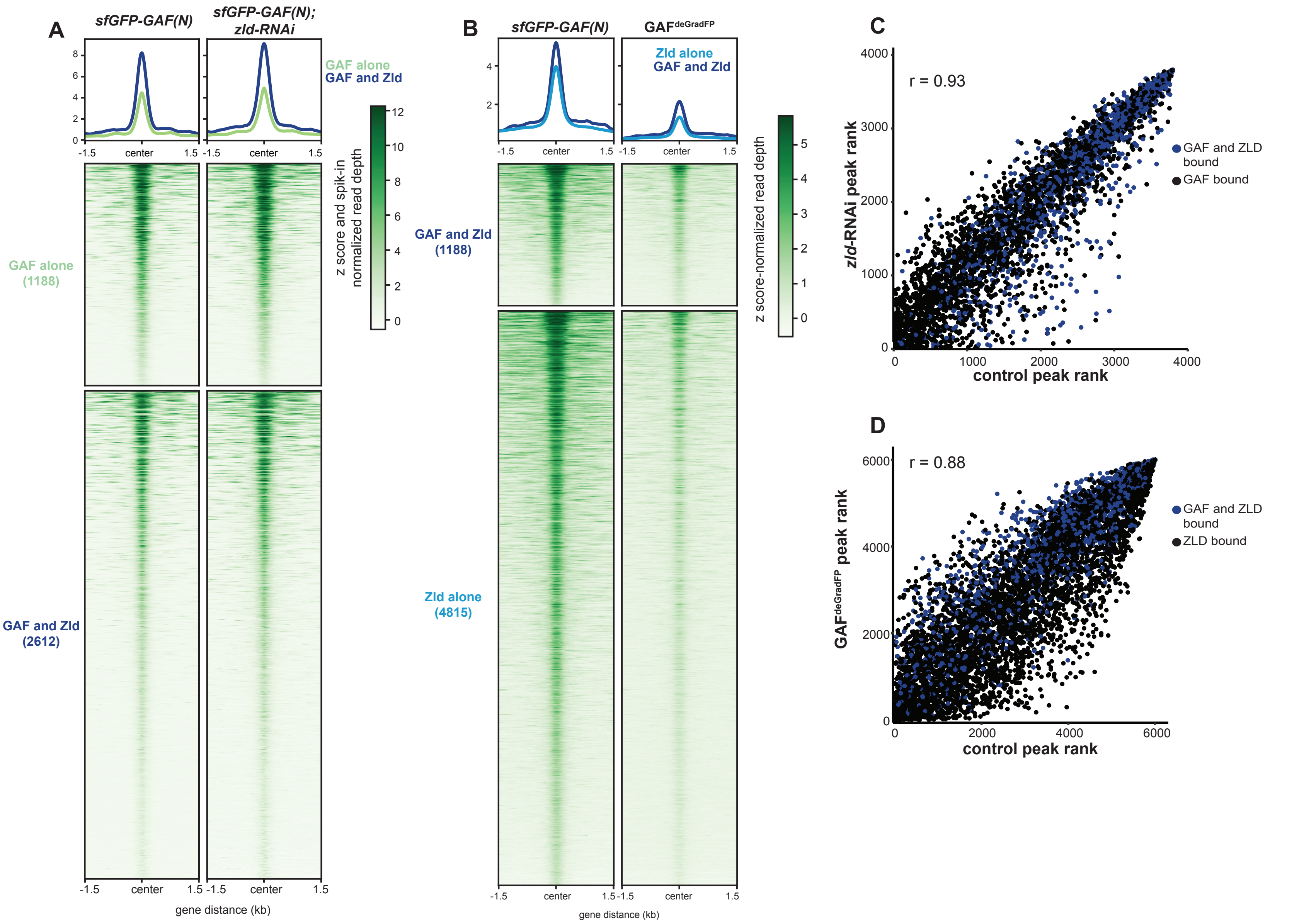


**Figure 4 - figure supplement 2: Independent chromatin binding by GAF and Zld**. A. Heatmap of high-confidence, anti-GFP ChIP-seq peaks from 2-2.5 hr AEL *sfGFP-GAF(N)* and *zld*-RNAi;*sfGFP-GAF(N)* embryos. The heatmap is divided into sites that have both GAF and Zld binding and where GAF binds independently of Zld. B. Heatmap of high-confidence, anti-Zld ChIP-seq peaks from *sfGFP-GAF(N)* control and GAF$^{deGradFP}$ embryos at 2-2.5 hr AEL. The heatmap is divided into sites that have both GAF and Zld binding and where Zld binds independently of GAF. C. Correlation between ranked peak heights of ChIP for GFP from (A). Peaks are ranked such that the highest peaks have the highest ranking. Color indicates those regions that are bound by both GAF and Zld. D. Correlation between ranked peak heights of ChIP for Zld from (C). Peaks are ranked such that the highest peaks have the highest ranking. Color indicates those regions that are bound by both GAF and Zld.
