## Supplemental Figure 5-1 for "GAF is essential for zygotic genome activation and chromatin accessibility in the early *Drosophila* embryo"

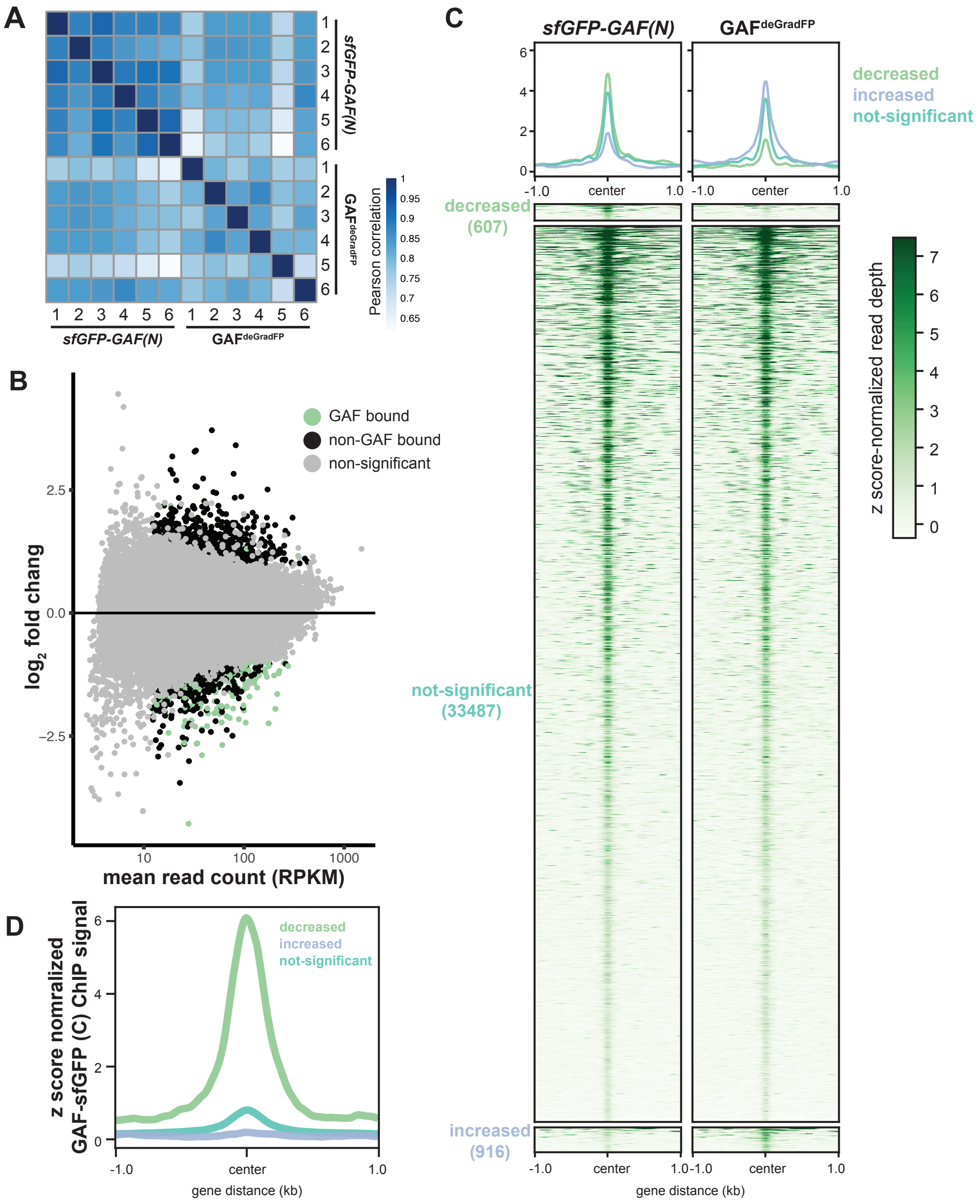


**Figure 5 - figure supplement 1: GAF is required for chromatin accessibility.** A. Heatmap of Pearson correlation coefficients between ATAC-seq replicates for GAF$^{deGradFP}$ and *sfGFP-GAF(N)* controls. B. MA plot of regions that change in accessbility in GAF$^{deGradFP}$ embryos as compared to *sfGPF-GAF(N)* controls. GAF binding was determined using GAF-sfGFP(C) ChIP-seq from stage 5 embryos. C. Heatmap of the ATAC-seq data for GAF$^{deGradFP}$ embryos and *sfGPF-GAF(N)* controls subdivided by whether the region increased, decreased or remained unchanged in accessibility in theGAF$^{deGradFP}$ embryos as compared the controls. D. Average GAF-sfGFP(C) ChIP-seq signal from stage 5 embryos for regions that increased, decreased or remained unchanged in accessbility in the GAF$^{deGradFP}$ embryos as compared to *sfGFP-GAF(N)* controls.
