## Supplemental Figure 6-1 for "GAF is essential for zygotic genome activation and chromatin accessibility in the early *Drosophila* embryo"

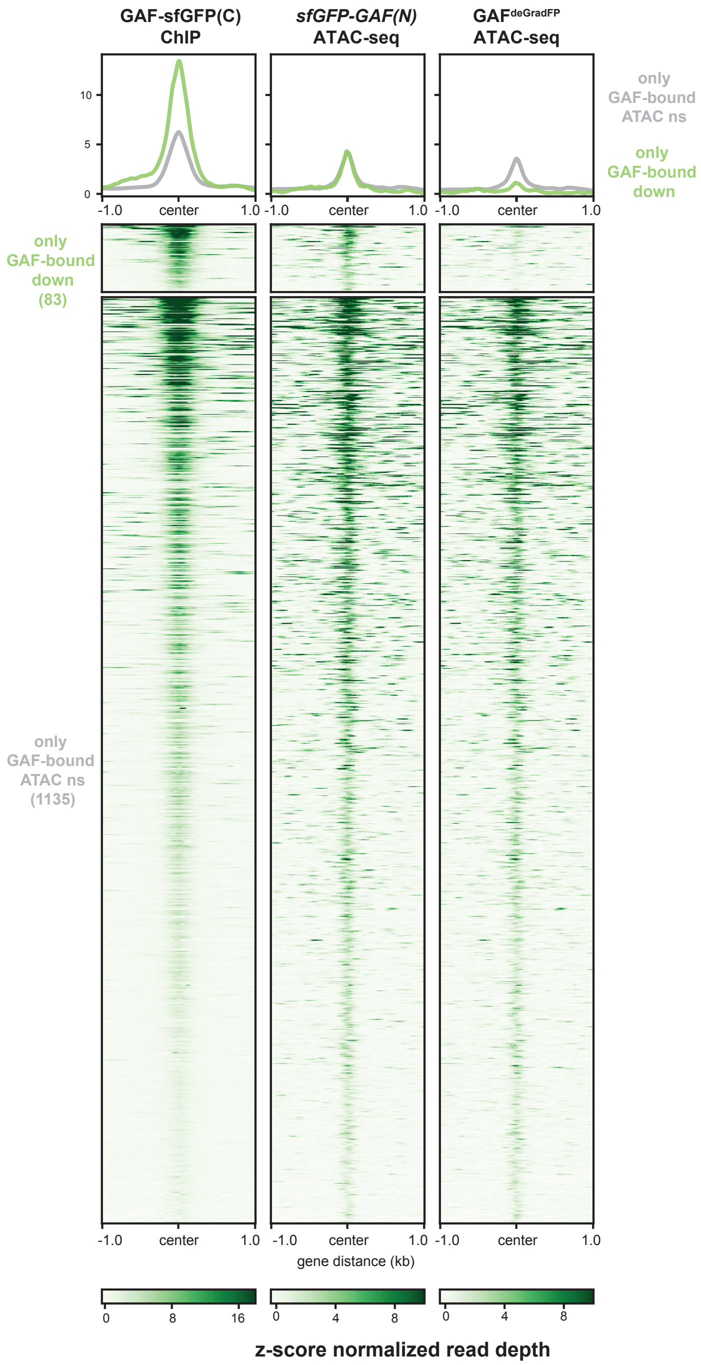


**Figure 6 - figure supplement 1: A subset of regions bound by GAF, and not Zld, depend on GAF for accessibility.** Heatmaps of regions that are bound by GAF (excluding GAF and Zld co-bound sites) as determined by *GAF-sfGFP(C)* stage 5 ChIP-seq data sub-divided by whether or not these regions change in accessibilty upon GAF depletion.
