## Supplemental Figure 6-2 for "GAF is essential for zygotic genome activation and chromatin accessibility in the early *Drosophila* embryo"

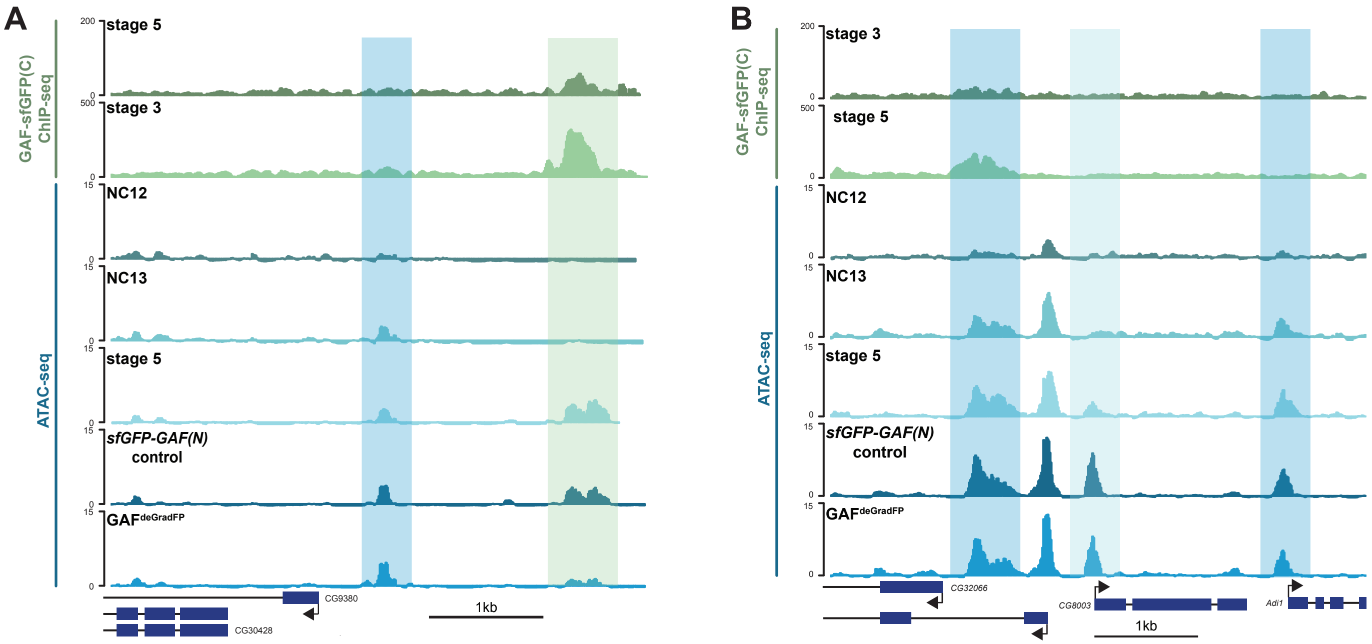


**Figure 6 - figure supplement 2: Regions that gain accessibility late during the MZT are accessible in $GAF^{deGraFP}$ embryos used for ATAC-seq.** A. Genome browser tracks showing a region that gains accessibility at NC13 that is maintained in $GAF^{deGradFP}$ embryos (dark-blue shading) and a region that gains accessibility at stage 5, is bound by GAF at stage 3 and stage 5, and at which accessibility is decreased in $GAF^{deGradFP}$ embryos (light-green shading). B. Genome browser tracks showing regions that gain accessibility at NC13 and are maintained in $GAF^{deGradFP}$ embryos (dark-blue shading) and a region that gains accessibility at stage 5 that is maintained in the $GAF^{deGradFP}$ embryos (light-blue shading).
